# Bursting deep dorsal horn neurons: the pharmacological target for the anti-spastic effects of Zolmitriptan?

**DOI:** 10.1101/075697

**Authors:** Eva Meier Carlsen, Rune Rasmussen

## Abstract

In a recent publication, Thaweerattanasinp and colleagues employed an in vitro preparation and electrophysiology to investigate firing properties of deep dorsal horn neurons following spinal cord injury during NMDA or zolmitriptan application. Deep dorsal horn neurons were classified into bursting, simple or tonic, with bursting neurons showing NMDA and zolmitriptan sensitivity. Here, we discuss the findings in a methodological framework and propose future experiments of importance for translating the results into a physiological setting.

## 1. INTRODUCTION

The ability to move in response to incoming external stimuli is likely a key-reason for why the process of evolution has resulted in the development of a central nervous system. All of our interactions with the surrounding world occurs on the foundation of our ability to generate movements. The spinal cord is absolutely essential for this generation of movement. In the ventral horn of the spinal cord, lower motoneurons send their axons directly to skeletal muscles, providing the final common pathway for conveying neuronal commands to the muscles.

In the spinal cord, incoming information from sensory afferents and descending commands from supra-spinal areas integrate, to shape the output firing behavior of individual motoneurons. In order to generate flexible yet reliable movements, it is crucial that the motoneuron assemblies receive the appropriate balance of excitatory and inhibitory synaptic barrages. Following spinal cord injury (SCI), this balance may be disturbed, frequently leading to involuntary motor activity and episodic spasticity in patients suffering from SCI (Adams and Hicks 2005; McKay et al. 2011).

Several monoamine neuromodulators are known for their potent ability to regulate spinal locomotor networks: one of them is 5-hydroxytryptamine (5-HT, or serotonin). In mammals, nearly all of the serotonin originates from dedicated neurons in the raphe nuclei, the primary source of serotonin in the central nervous system. Axons located in the lower raphe nuclei project diffusely into all spinal cord segments, providing strong serotonergic modulation of locomotor networks. The raphe-spinal pathway targets many types of neurons and receptors. Activation of the G_i_-protein-coupled 5-HT_1B/1D/1F_ is suggested to facilitate inhibition and decrease neuronal excitability of dorsal horn neurons (Yoshimura and Furue 2006). However, the serotonin receptor system is immensely rich in diversity, and thus the composition of serotonin receptor activation, at a given time, can determine if serotonin overall increases or decreases motoneuron excitability (Cotel et al. 2013).

As a consequence of SCI, axonal fibers connecting the raphe-spinal pathway may be transected, leading to a loss of serotonergic innervation to all spinal segments below the lesion (Murray et al. 2010). Conversely, glutamate, the main excitatory transmitter, is intrinsically produced in neurons located within the spinal cord, and is thus still present following SCI. Therefore, chronic SCI will likely be accompanied by a shift in glutamatergic to serotonergic transmission ratio in spinal locomotor networks. This loss of raphe-spinal-mediated serotonergic transmission, leading to reduced 5-HT_1B/1D/1F_ receptor activation, may consequently lead to a loss of inhibitory tone, and eventually push the motoneurons into a hyper-excitable state (Hains et al. 2003). In such a hyper-excitable state, glutamate released from spinal interneurons, could, ultimately, elicit muscle spasms. Indeed, involuntary motor activity and spasticity is a frequent symptom of chronic SCI, and in in vitro models of SCI, dorsal root stimulation evokes unusually long NMDA receptor-dependent excitatory postsynaptic potentials (Bennett et al. 2004).

Zolmitriptan, a 5-HT_1B/1D_ receptor agonist can alleviate muscle spasms in vivo and reduce the neuronal hyper-excitability in response to dorsal root stimulation in vitro (Murray et al. 2011), but it is currently unknown where, and what cell types, in the spinal cord, the drug exerts its beneficial actions. One suggested target is the glutamatergic spinal deep dorsal horn (DDH) neurons, receiving synaptic input from group I and II sensory afferents (Murray et al. 2011), well-positioned for providing excessive excitatory barrages onto motoneurons. In a recent article published in *Journal of Neurophysiology*, Thaweerattanasinp and colleagues (2016) therefore set out to investigate the firing properties of DDH neurons and unravel their response patterns to NMDA and zolmitriptan application following SCI.

## 2. OVERALL FINDINGS

In this study, DDH neurons were classified into one of three groups: burst, simple or tonic firing. This classification was based on single-neuron firing activity, assessed with extra-cellular recordings made with sharp glass microelectrodes in vitro. Thaweerattanasinp and colleagues show, that burst firing DDH neurons respond to NMDA application with an increase in burst duration and spontaneous firing rate, and to zolmitriptan with a decrease in burst duration, evoked spike count and spontaneous firing rate. Simple firing DDH neurons did not significantly change any of their firing properties in response to neither NMDA nor zolmitriptan application. Unfortunately, the authors were only able to record from a limited number of tonic neurons during drug application and their response patterns could thus not be evaluated in a quantitatively satisfying manner.

## 3. DETAILED DISCUSSION AND FUTURE EXPERIMENTS

All of the present experiments were carried out in a model of acute spinal transection using the in vitro sacral cord preparation. In this model, the authors of the study argues, that NMDA bath application mimics the scenario after SCI where intrinsically-produced glutamate, in the absence of descending serotonergic release, triggers excessive bursting of DDH neurons, ultimately leading to muscle spasms. On the contrary, zolmitriptan could potentially dampen this exaggerated burst firing by activating serotonergic G_i_-coupled 5-HT_1B/1D_ receptors on DDH neurons, consequently reducing synaptic barrages onto motoneurons and alleviating muscle spasms. Taking a closer look at the heterogeneous bursting activity of control DDH neurons (measured by *e.g.* evoked spike count and burst duration) in Figure 3 and 4, it is, however, intriguing that, in the experiments with NMDA bath application, the control group is represented by low burst duration neurons, whereas, for the zolmitriptan experiment, the neurons in the control group has a high burst duration. From this observation, a complementary interpretation of the results from Thaweerattanasinp and colleagues might be, that NMDA and zolmitriptan, not only affects DDH neurons in a pathological setting after SCI, but also act as opposing modulators of DDH neuron burst duration within their physiological spectrum. In this view, activation of 5-HT_1B/1D_ receptors with zolmitriptan do not have a general inhibitory effect, but only suppresses the high burst duration neurons and pulls them back into the intermediate zone of the burst duration spectrum. This view, would also state that NMDA do not necessarily cause exacerbated firing of DDH neurons, but rather is a modulator that pushes low burst duration neurons into the intermediate zone of the burst duration spectrum. The interpretation, that NMDA application normalizes the burst duration of low burst duration neurons also allows for the puzzling finding, that Thaweerattanasinp and colleagues measured no concurrent increase in evoked spike count of the DDH neurons following NMDA application. The latter observation may appear somewhat incompatible with the rationale, that glutamate, in the absence of serotonin, facilitates excessive bursting of DDH neurons, eliciting increased excitatory synaptic barrages onto motoneurons, ultimately eliciting muscle spasm occurrence.

Another potential source for the lack of NMDA-mediated effect on the number of evoked spikes might be inherent to the computational analysis procedure chosen by the authors. For determining the number of evoked spikes a 1-second time frame following the sensory stimulus onset was used, whereas, burst duration was defined as the temporal interval from the first to the last evoked spike within this 1-second time frame. For the bursting DDH neurons, the evoked spike count was subsequently corrected for the spontaneous firing rate before the stimulus onset, whereas this was not plausible for the burst duration measure. As can be observed from Figure 3, NMDA application triggered a significantly increased spontaneous firing rate in the ~ 1 to 2 Hz spectra. Consequently, determining burst duration as the interval between the first and last spike occurrence, in a 1-second time frame, is likely to be contaminated with one or maybe two spontaneous spikes, thus broadening the burst duration, whereas the evoked spike count, given the correction-method, will be unsusceptible to this effect. This complication might possibly have produced longer burst durations, but maybe even more importantly, may also have masked slight NMDA-mediated effects on the number of evoked spikes. One possible remedy could be to determine the evoked spike count in a temporally narrower time frame, *e.g.* 20-50 milliseconds following stimulus onset, thereby reducing the likelihood of having spontaneous spikes contained within the evoked bursting period. If a narrower time frame reveals increased numbers of evoked spikes in the immediate burst following stimulus onset, this would support the rational that NMDA receptor activation do indeed facilitate firing of bursting DDH neurons, and thus that NMDA-induced bursting of these neurons could contribute to the generation of muscle spasm after SCI.

Thaweerattanasinp and colleagues report that they used three different NMDA concentrations for unraveling NMDA receptor-mediated effects, ranging from 15 to 100 *µ*M, and given that the authors made no apparent distinction between these, relatively different, concentrations in their result section, we assume all concentrations produced comparable effects. This assumption, together with the subtle changes observed in firing properties of bursting DDH neurons might indicate, that NMDA receptor-mediated processes were already close to saturated within the spinal cord preparation used by the authors. To evaluate the validity of this interpretation, one purposeful experiment would be to apply an NMDA receptor antagonist, such as APV (2-Amino-5-phosphonopentanoic acid), and determine the firing properties of bursting DDH neurons in the spinal cord preparation, without prior exogenous NMDA bath application. Overall for the NMDA application experiments, considering that dorsal root stimulation evoked spiking of bursting DDH neurons did not increase following NMDA application, the effect of glutamate and NMDA receptor activation, in the context of excessive DDH neuron firing, facilitating muscle spasms in chronic SCI, does not appear trivial and linear. Furthermore, given that the present experiments were performed in an acute in vitro SCI preparation, it may not be fully appropriate to draw bold conclusions and extrapolate these results to a chronic in vivo SCI phenotype.

The 5-HT_1B/1D_ receptor agonist zolmitriptan alleviates muscle spasm symptoms following chronic SCI (Murray et al. 2011). Here, Thaweerattanasinp and colleagues show, that zolmitriptan suppresses the firing properties of bursting DDH neurons following acute SCI. This overall effect was manifested in decreased evoked spike count, shortened burst duration and lowered spontaneous firing rate. Thus, pointing to bursting DDH neurons as a pharmacological target for the anti-spastic effects of zolmitriptan.

Lower motoneurons provide the final common pathway for conveying neuronal commands to the muscles. Therefore, investigating if zolmitriptan lowers the excitability of motoneurons, correlated to suppression of bursting DDH neurons, would be powerful, and could provide a mechanistic link between DDH neuronal bursting and changes in motoneuron excitability and firing following SCI. To experimentally interrogate this link, one approach could be to record from the ventral roots, comprising motoneuron bundles, while stimulating dorsal root sensory afferents and simultaneously record the activity of identified bursting DDH neurons. If zolmitriptan alleviate chronic SCI-induced muscle spasms, via DDH neuron modulation, one might expect that suppressing bursting activity of these neurons closely correlate with decreased excitability of motoneurons. Opposite, if these two measures correlate poorly, one explanation could be that the relationship between DDH neurons and motoneurons is not the same in an acute SCI preparation, that more mimics a state where motoneurons are virtually non-excitable, compared to the chronic SCI where motoneurons become hyper-excitable.

The comprehensive effect of the loss of serotonin for locomotor spinal networks is complex and somewhat unpredictable due to the immense diversity and differential expression of serotonin receptors on spinal neurons. In the chronic phase of SCI, spinal motoneurons become super-sensitive to serotonin (Barbeau and Bdard 1981), plausibly due to dramatic up-regulation of constitutively active G_q_-coupled 5-HT_2A/2C_ receptors (Ren et al. 2013). By facilitating persistent inward currents, serotonin, via 5-HT_2A/2C_ receptor activation, may chronically lower the firing threshold of spinal motoneurons, suggested as an important step for the etiology of abnormal muscle activity and spasms following SCI.

To investigate the impact of zolmitriptan-mediated suppression of bursting DDH neurons, on motoneuron excitability, in the chronic SCI, one approach could be to mimic the constitutive 5-HT_2A/2C_ receptor activation by bath applying appropriate agonists for these receptors. This, in theory, would push the spinal motoneurons into the hyper-excitable state observed in chronic SCI, by lowering their firing threshold. Thus, by stimulating sensory afferents and recording from ventral root motoneurons in 5-HT_2A/2C_ receptor agonist-containing media, with or without zolmitriptan, one could gain insight to the effects of zolmitriptan in the chronic SCI phenotype. Result from the depicted experiment could thereby expand the clinical relevance and impact of the present findings made by Thaweerattanasinp and colleagues.

## 4. CONCLUSION

Taken together, Thaweerattanasinp and colleagues reports the existence of three types of DDH neurons; burst, simple and tonic based on their firing properties. The bursting DDH neurons were suppressed by the 5-HT_1B/1D_ receptor agonist zolmitriptan, known to alleviate muscle spasms in chronic SCI. NMDA application did modulate the bursting neurons as well, but the results are less clear and methodological changes and further experiments are recommended for significant conclusions to be made. It should be noted that, especially the bursting DDH neurons appear quite heterogeneous, especially with respect to spontaneous firing frequencies. As the spontaneous firing rate was affected by both NMDA and zolmitriptan, a further sub-division into tonic and non-tonic bursting DDH neurons could be appropriate and might reveal unexpected differences in other firing properties. Treatment with zolmitriptan may cause unwanted side effects; since it activates all 5-HT_1B/1D_ receptors found on all types of spinal neurons. If the DDH neurons truly are the mediator of this drugs anti-spastic effects, understanding the mechanism behind could form the foundation for the development of a more specific drug. In summary, by employing a model of acute SCI in conjunction with electrophysiology and pharmacology, the work by Thaweerattanasinp paves the way for future research aiming at investigating the detailed neurophysiological mechanisms underlying the positive clinical effects of zolmitriptan, as well as for interrogating the role of 5-HT_1B/1D_ receptors in spinal locomotor networks.

## 6. AUTHOR CONTRIBUTIONS

E.M.C. and R.R. drafted manuscript; E.M.C. and R.R. edited and revised manuscript; E.M.C. and R.R. approved final version of manuscript.

